# Computational prediction of structure, function and interaction of aphid salivary effector proteins

**DOI:** 10.1101/2023.10.02.560451

**Authors:** Thomas Waksman, Edmund Astin, S. Ronan Fisher, William N. Hunter, Jorunn I.B. Bos

## Abstract

Similar to plant pathogens, phloem-feeding insects such as aphids deliver effector proteins inside their hosts that act to promote host susceptibility and enable feeding and infestation. Despite exciting progress towards identifying and characterizing effector proteins from these insects, their functions remain largely unknown. The recent ground-breaking development in protein structure prediction algorithms combined with the availability of proteomics and transcriptomic datasets for agriculturally important pests, such as the aphid *Myzus persicae* (green peach aphid), provides new opportunities to explore the structural and functional diversity of effector repertoires. In this study, we sought to gain insight into the the *M. persicae* effector repertoire by predicting and analysing the structures of a set of 71 effector candidate proteins. We used two protein structure prediction methods, AlphaFold and OmegaFold, which produced mutually consistent results. We observed a wide continuous spectrum of sizes and structures among the effector candidates, from disordered proteins to globular enzymes. We made use of the structural information and state-of-the-art computational methods to predict *M. persicae* effector protein properties, including function and interaction with host plant proteins. Overall, our investigation provides novel insights into the structure, function, and interaction prediction of aphid effector repertoires and will guide the necessary experimental characterization to address new hypotheses.

## Introduction

Like other plant parasites, phloem-feeding insects, belonging to the order Hemiptera, deliver diverse effector proteins from their saliva inside the host plant during the infestation process to modulate host responses (reviewed by Naalden *et al*., 2021; Wang *et al*., 2023). Among these insects are major agricultural pests, including aphids, which are difficult to control due to the lack of genetic crop resistances and reliance on pesticides, which are damaging to the environment and lead to evolution of pesticide resistance. Over the past decade the number of aphid species with available genome and/or transcriptome data has increased rapidly in combination with saliva proteomic approaches, this has facilitated the identification of candidate effector repertoires from various aphid and Hemipteran species (reviewed by Naalden *et al*., 2021; Wang *et al*., 2023). Characterisation of salivary effector repertoires will provide new insights into how insects are able to infest plants and inform plant protection strategies which interfere with effector protein action.

*Myzus persicae* (green peach aphid), is a major pest of crop plants globally, in part due to its exceptionally broad host range and ability to vector many important plant viruses (Blackman & Eastop, 2000). The effector repertoire of *M. persicae* consists of diverse proteins, as predicted by saliva proteomics and bioinformatics pipelines based on either salivary gland or head-versus-body transcriptomic data (Bos *et al*., 2010; Harmel *et al*., 2008; Thorpe *et al*., 2016; Thorpe *et al*., 2018; Vandermoten *et al*., 2014). Whilst many of these proteins are present across aphid species with differing levels of sequence diversity, there is also evidence of potentially aphid species-specific effectors (Boulain *et al*., 2018; Thorpe *et al*., 2016; Thorpe *et al*., 2018). For some aphid effectors/effector families there is evidence of gene duplication and diversification, pointing to potential neofunctionalization (Boulain *et al*., 2018; Dommel *et al*., 2020; Thorpe *et al*., 2018). Functional annotation of predicted aphid effectors that are conserved across insect species based on sequence similarity searches point to roles in enzymatic activities (reviewed by Naalden *et al*., 2021; Wang *et al*., 2023). Common enzymatic activities among predicted effectors are in carbohydrate metabolism and redox activity, which may be important for feeding/digestion, weakening of the plant cell wall, and suppression of plant defenses. However, sequence-based similarity searches for the majority of predicted aphid effectors, especially those with less sequence conservation, do not point to any potential functions. Despite the unknown activity of these more divergent effectors, some of these proteins are known to target host plant proteins to achieve desired manipulation of host processes (Chaudhary *et al*., 2019; Liu *et al*., 2022; Rodriguez *et al*., 2017). Importantly, effectors from different kingdoms of life may share common host target proteins, suggesting convergent evolution of virulence strategies (Liu *et al*., 2022; Lozano-Torres *et al*., 2012; Mukhtar *et al*., 2011; Weßling *et al*., 2014). However, despite progress in aphid effector identification, our knowledge on how these proteins alter the host environment to promote infestation remains limited.

While there is minimal knowledge of the structure of herbivorous insect effector proteins, structural biology of plant fungal pathogen effectors has provided new insights into mechanisms underlying effector evolution (Franceschetti *et al*., 2017; Outram *et al*., 2022). Initially, structural homology among effectors was experimentally determined and identified novel folds in effectors belonging to the WY family (Boutemy *et al*., 2011; He *et al*., 2019), RNase-like proteins expressed in Haustoria (RALPH) family (Bauer *et al*., 2021; Spanu, 2017), *Magnaporthe* Avrs and ToxB-like (MAX) family (de Guillen *et al*., 2015), *Leptosphaeria* AviRulence and Suppressing (LARS) family (Lazar *et al*., 2022), and ToxA family (Di *et al*., 2017). Recent impressive developments in algorithms for protein structure prediction and structure comparison now allow large-scale structural bioinformatics investigations into protein evolution, which make use of the higher conservation of protein structure compared to sequence (Jumper *et al*., 2021; van Kempen *et al*., 2023). For example in plant pathogenic fungal species, structural bioinformatics of whole secretomes unveiled new effector families and detected common ancestry between proteins which have greatly diverged in sequence, especially at surface-exposed residues (Derbyshire & Raffaele, 2023; Seong & Krasileva, 2021; Seong & Krasileva, 2023).

Computational analyses of phytopathogen effector structures also showed that regions of intrinsic disorder are common among effector proteins, with some proteins featuring a combination of disordered/ordered regions and others with disordered regions spanning most of the primary sequence (Derbyshire & Raffaele, 2023; Marin *et al*., 2013; Seong & Krasileva, 2023; Yu *et al*., 2023). Intrinsic disorder among pathogen effectors could be linked to mechanisms of effector translocation, evasion of immunity, as well as dynamic host protein targeting (Marin *et al*., 2013), and the presence of disordered regions may drive effector diversification through domain fusions (Seong & Krasileva, 2023). With intrinsically disordered regions being prevalent in pathogen effectors, there is a need to explore their functions, both in triggering immunity as well as in the potential dynamic targeting of relevant proteins and biological processes in the host plant environment.

In this study, we apply structural bioinformatic methods to characterize the salivary effector protein repertoire from *M. persicae* (Fig. 1). We use two stateof-the-art protein structure prediction algorithms, and the structure information as input for algorithms to predict protein function and protein interaction. Notably, we do not exclude disordered proteins which lack a precise structure prediction from analysis: we prefer to acknowledge that these under-studied and unpredictable proteins are important agents of virulence which should be investigated through computational or experimental approaches. In future work, experimental determination of *M. persicae* effector structures and mechanisms of action will provide an improved understanding of insect herbivory, eukaryotic effector protein evolution and protein-protein interactions across species.

**Figure 1:**
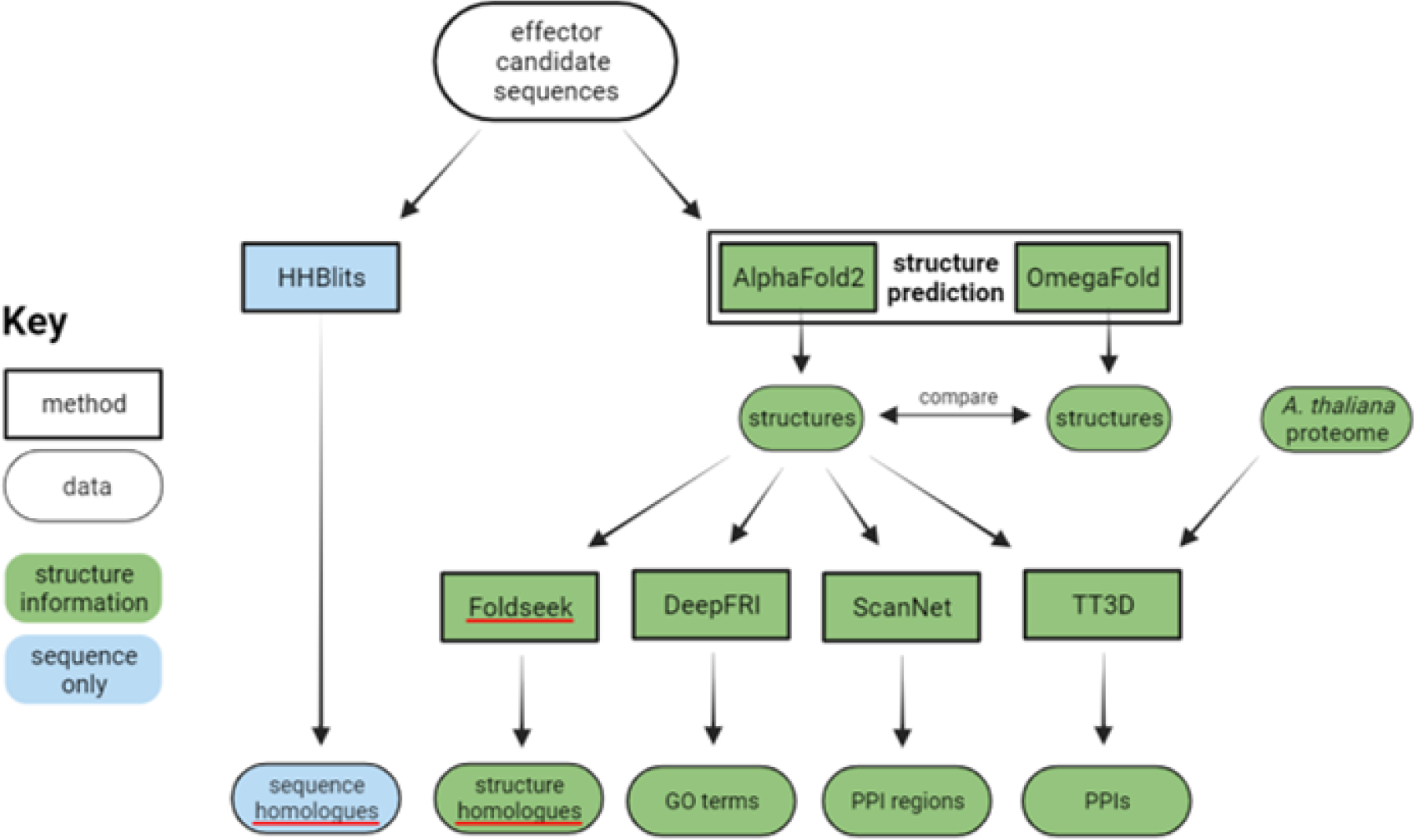
Design of the bioinformatics pipeline used to characterize *M. persicae* salivary effector candidate proteins. Protein sequences curated from published experimental studies were used for structure prediction by AlphaFold and OmegaFold. For each protein, structures predicted by AlphaFold and OmegaFold were compared and AlphaFold structures were used for prediction of functional and interactive properties. DeepFRI was used to predict Gene Ontology (GO) terms, ScanNet was used to predict amino acids involved in protein interaction and Topsy Turvy 3D (TT3D) was used to predict *A. thaliana* proteins which interact with the effector candidates. Foldseek and HHBlits were used for structure-based and sequence-based database searching respectively, to annotate function based on homology. Green color indicates the parts of the pipeline that involve protein structure information. Created in BioRender.com.

## Results

### Structure prediction reveals great diversity of proteins in the *M. persicae* salivary effector repertoire

We conducted a literature review to construct a list of *M. persicae* candidate effector protein sequences. These gene products have been detected in secreted saliva by proteomics, or salivary glands by transcriptomics (Bos *et al*., 2010; Elzinga *et al*., 2014; Harmel *et al*., 2008; Liu *et al*., 2022; Thorpe *et al*., 2016; Vandermoten *et al*., 2014). The proteins, including previously characterized effectors, were each named ‘Mp’ concatenated with a unique identification code, up to Mp95 (Supplementary File 1). Sequences with any of the following characteristics were removed: incomplete, negative for signal peptide, positive for transmembrane domain, or length >700. The structures of the 71 remaining proteins (with signal peptide removed) were predicted using AlphaFold (AlphaFold 2) as well as OmegaFold via ColabFold (Jumper *et al*., 2021; Mirdita *et al*., 2022; Ruidong *et al*., 2022). These algorithms are reported to achieve the highest accuracy available among many competing protein structure prediction tools, yet differ significantly in their method. AlphaFold is primarily trained on the Protein Data Bank (PDB) (wwPDB Consortium, 2019) and requires a multiple sequence alignment to predict the protein structure, whereas OmegaFold is a language model trained on UniRef50 protein sequences and does not construct a multiple sequence alignment to predict the structure (Jumper *et al*., 2021; Ruidong *et al*., 2022).

Structure prediction indicated that the *M. persicae* effector set contains a wide range of structures, with varying degrees of disorder or globularity (Fig. 2). The degree of disorder is quantified using the ratio of the protein’s solvent-accessible surface area (SASA) to length, which shows strong negative correlation with confidence of the structure prediction (Supplementary Fig. 1A). Numerous proteins with high SASA/length ratio and low prediction confidence are likely to be intrinsically disordered proteins which do not exist in a fixed conformation, and the structure prediction may be considered as an accurate depiction of one possible conformation of the protein. AlphaFold and OmegaFold predict a similar degree of disorder for each protein and nearly identical structures for proteins with a well-defined tertiary structure, indicated by very high DALI Z score (Supplementary Fig. 1B). Furthermore, the profile of per-residue confidence scores along the length of each protein generally shows a strong positive correlation between AlphaFold and OmegaFold (Supplementary Fig. 1C). As these two structure prediction algorithms are fundamentally different, their close agreement may lead to much higher confidence in the accuracy of the protein structure predictions, compared to using a single structure prediction method.

**Figure 2:**
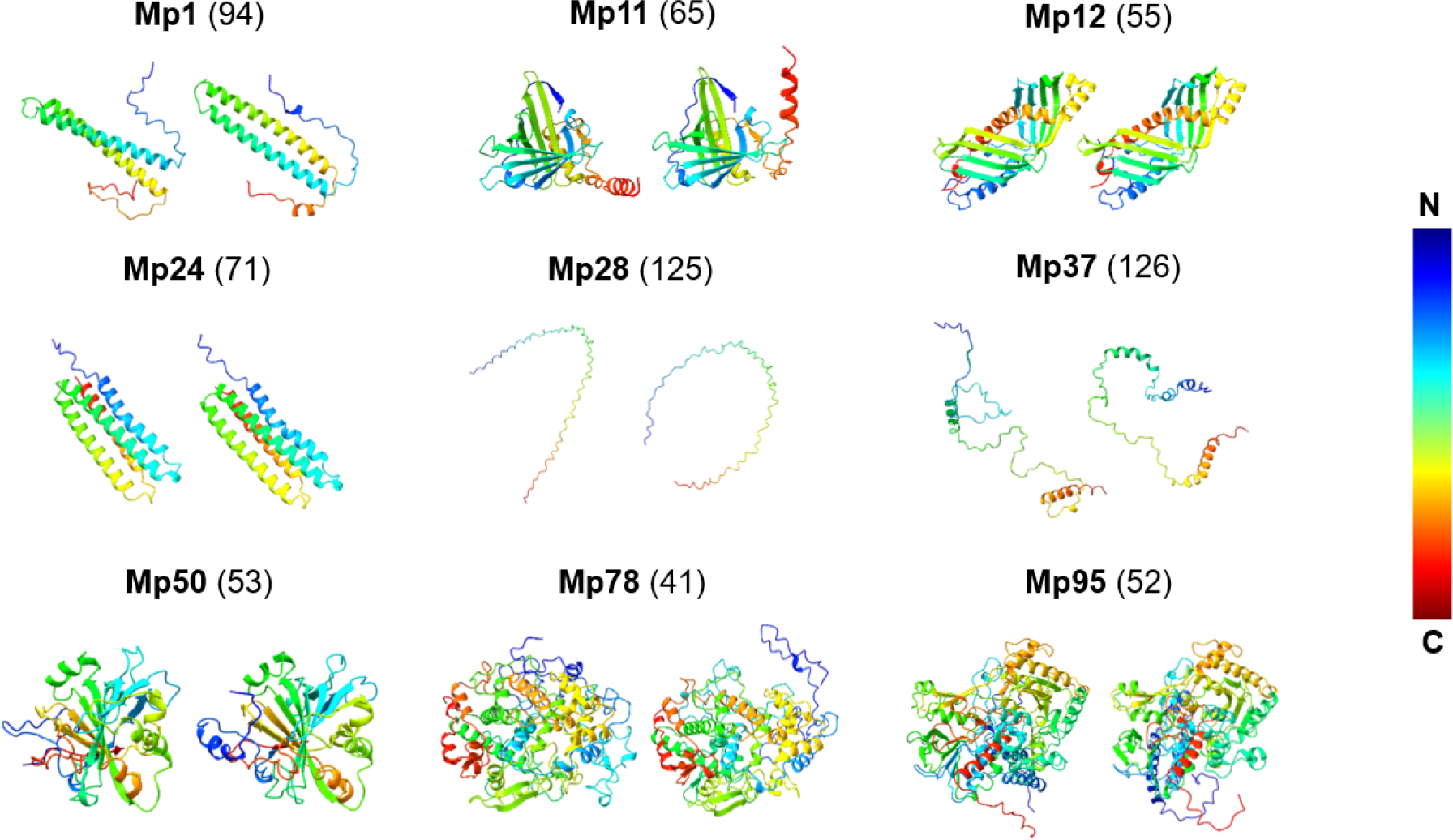
Examples of *M. persicae* salivary effector candidate predicted protein structures. For each protein, the AlphaFold and OmegaFold predictions are shown on left and right, respectively. Each structure is colored as a spectrum from N-terminus to C-terminus, as indicated by the color bar. The solvent accessible surface area (SASA)/length ratio (Å2 per amino acid) of the AlphaFold prediction is indicated in parentheses.

AlphaFold was also used to predict the structure for an equally sized dataset of random *M. persicae* proteins (Supplementary File 1). The random set and the effector candidate set have a similar average SASA/length ratio, suggesting that the degree of disorder of *M. persicae* salivary effectors is similar to *M. persicae* proteins in general, while the average SASA/length ratio of an equally sized dataset of random proteins from the PDB is much lower (Supplementary Fig. 1D). These results emphasize that there are numerous *M. persicae* salivary effector proteins of interest which lack similarity to proteins in the PDB these proteins may not be amenable to structure determination using conventional experimental methods, or structure prediction methods that are trained on the PDB. From now on we refer to the half of the effector candidates with lowest SASA/length ratio as ‘ordered’ and the half with highest SASA/length ratio as ‘disordered’. In general, the proteins in the ‘ordered’ half of the dataset are relatively globular with well-defined tertiary structure, while those in the ‘disordered’ half lack this structure.

To investigate relationships between the effector candidates, pairwise comparison among all the proteins was performed based on sequence or structure, followed by clustering to group similar proteins together. Generally, similar clusters were observed based on sequence or structure, indicating that similarity between the proteins is usually of both types (Supplementary Fig. 2). However, similarity between some pairs of proteins could be detected based on either sequence or structure but not both (Supplementary Fig. 2). Among the proteins with clear tertiary structure, there are numerous different folds (supplementary data).

### Function prediction indicates protein activities enriched in the *M. persicae* salivary effector repertoire

After structure prediction, we aimed to predict functions of the *M. persicae* salivary effector candidates, combining structure and sequence information. Two methods were used: deep learning trained to predict functional information and searching for annotated homologous proteins in databases. For the former method, we used DeepFRI, which predicts Gene Ontology (GO) terms and requires the protein structure for best performance (Gligorijević *et al*., 2021); for the latter method, we used HHBlits for database searching based on sequence or Foldseek for database searching based on structure (Remmert *et al*., 2011; Steinegger *et al*., 2019; van Kempen *et al*., 2023). Foldseek represents a major advance in capability to discover homologous proteins, being efficient enough to quickly search large databases, and capable of local structure alignment of sub-sections of proteins. This enables detection of distant evolutionary relationships where structure is more conserved than sequence and highlights examples of effector evolution in which globular structurally conserved regions have fused with variable disordered regions (Barrio-Hernandez *et al*., 2023; Illerg å rd *et al*., 2009; Seong & Krasileva, 2023).

DeepFRI predicted GO terms enriched in the *M. persicae* effector set were discovered by comparison to the random *M. persicae* protein set. The effector candidates with relatively ordered structure are enriched for GO terms associated with cell signalling and enzyme activities such as carbohydrate degradation; the relatively disordered proteins are enriched for GO terms associated with protein and host cell interaction (Fig. 3).

**Figure 3:**
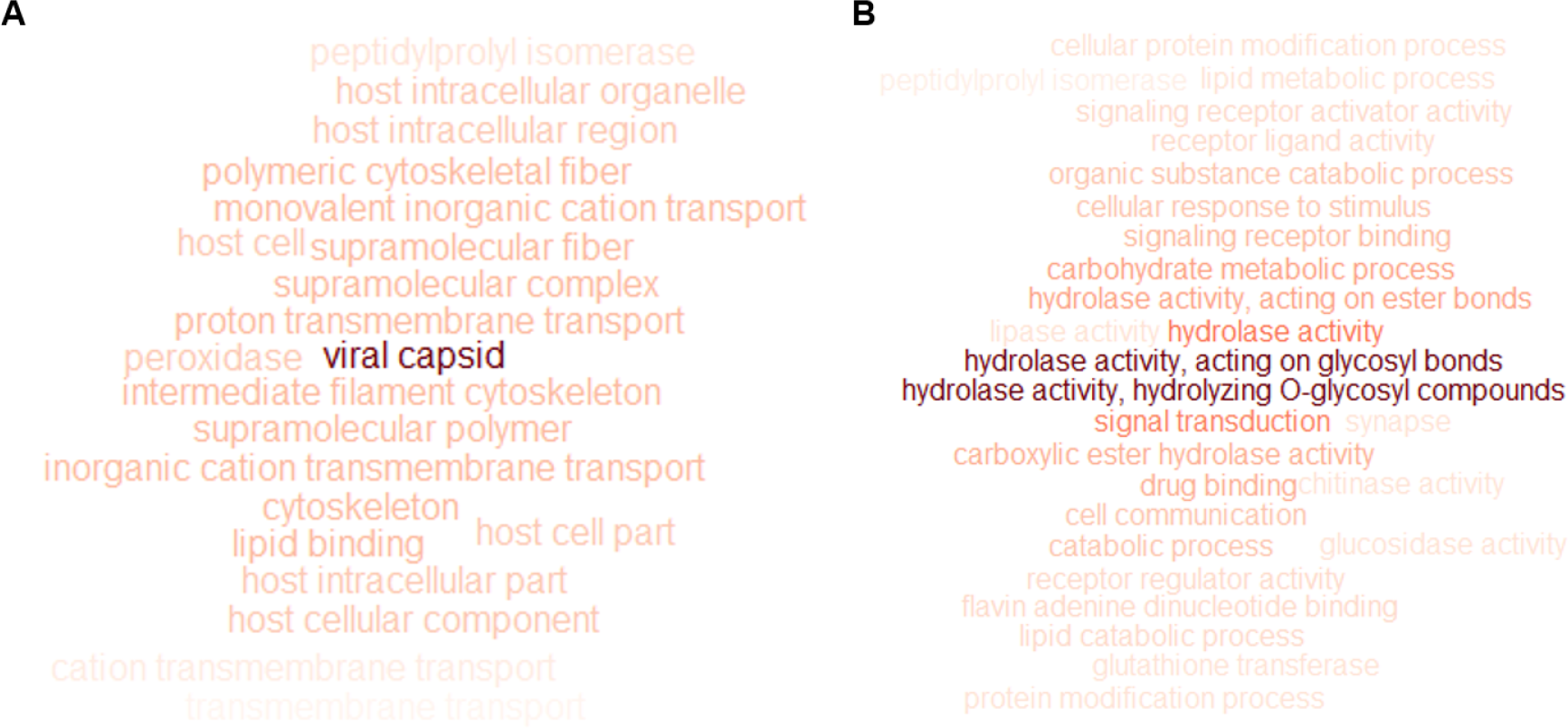
Gene Ontology (GO) terms enriched in the *M. persicae* salivary effector candidate dataset compared to the M. persicae random protein dataset, for **A**, the 36 effector candidates with highest solvent accessible surface area (SASA)/length ratio and **B**, the 36 effector candidates with lowest SASA/length ratio. DeepFRI-predicted GO terms were counted and weighted by confidence score; GO terms relating to protein localization and nucleotide metabolism were excluded. Visualization was performed using the R wordcloud package; GO term text color darkness is positively correlated with enrichment.

For many ordered proteins, searching databases for annotated homologues resulted in high confidence in predicted protein functions; for many of the disordered proteins, few strong homologous matches were returned, and the list consisted predominantly of poorly annotated proteins (supplementary data). Database searching by sequence rather than structure was generally more powerful to find annotated homologues for disordered proteins, which is expected because the sequence must be more conserved than the structure if the structure represents only one of many possible conformations. However, for some proteins structure-based searching with Foldseek recovered more significant matches than sequence-based searching with HHBlits. This was the case for three proteins which are associated with functional data (Mp1, Mp32, Mp58), demonstrating the expected benefit of using structure to detect homology (supplementary data). Overall, most of the larger effector candidates could be confidently annotated as enzymes of various types, and numerous proteins were annotated with a function related to hydrophobic small molecule binding (Supplementary File 1). Some proteins were annotated with a function suggesting a role within the insect rather than effector, including hormonal (e.g. bursicon, protein quiver, cardioactive peptide) or structural (e.g. cuticle protein) functions these proteins might be contaminants from tissue other than saliva in proteomics experiments, or secreted from salivary glands for function other than effector (Supplementary File 1). We noted that proteins annotated as cuticle proteins are responsible for the ‘viral capsid’ enriched GO term among the disordered effector candidates (Fig. 3A, Supplementary File 1, supplementary data) DeepFRI appears to have recognised a structural or invasive function for these proteins, although they do not originate from viruses. Approximately 30 out of 71 of the effector candidates remain without annotated function.

### Interaction prediction suggests mechanisms of *M. persicae* salivary effector evolution and function

Effector proteins are expected to interact with host target proteins to induce perturbation of the host to the benefit of the pest or pathogen. Therefore, we attempted to predict the plant proteins which may be targeted by the *M. persicae* effector repertoire, and how the effector proteins may interact with targets. ScanNet is a recently developed deep learning model achieving substantial success in predicting sites of protein interaction (Tubiana *et al*., 2022). Analysing the *M. persicae* effector candidates, disordered proteins were assigned a higher probability of being involved in protein interactions, compared to ordered proteins (Fig. 4A). For disordered proteins, higher probability of interaction was predicted for hydrophobic amino acids compared to other amino acid types, and this was not the case for ordered proteins (Fig. 4B). Hydrophobic residues would normally be found in the interior of a folded protein, as solvent (water) interaction is thermodynamically unfavourable. For disordered effector proteins, these hydrophobic patches may be their preferred sites of interaction with host targets, as the favourable solvent enthalpic effect upon binding would compensate for the unfavourable entropic effect of fixing the conformation of the protein. ScanNet consistently assigned an extremely low probability of interaction to the internal residues of the globular enzymes, distinguishing these hydrophobic amino acids that drive protein folding from hydrophobic amino acids of disordered proteins. However, several of these enzymes contain regions on the periphery or separate to the globular folded region, which are assigned a high probability of interaction (Fig. 4C). These regions may have evolved to confer effector activity, separate to the enzymatic activity.

**Figure 4:**
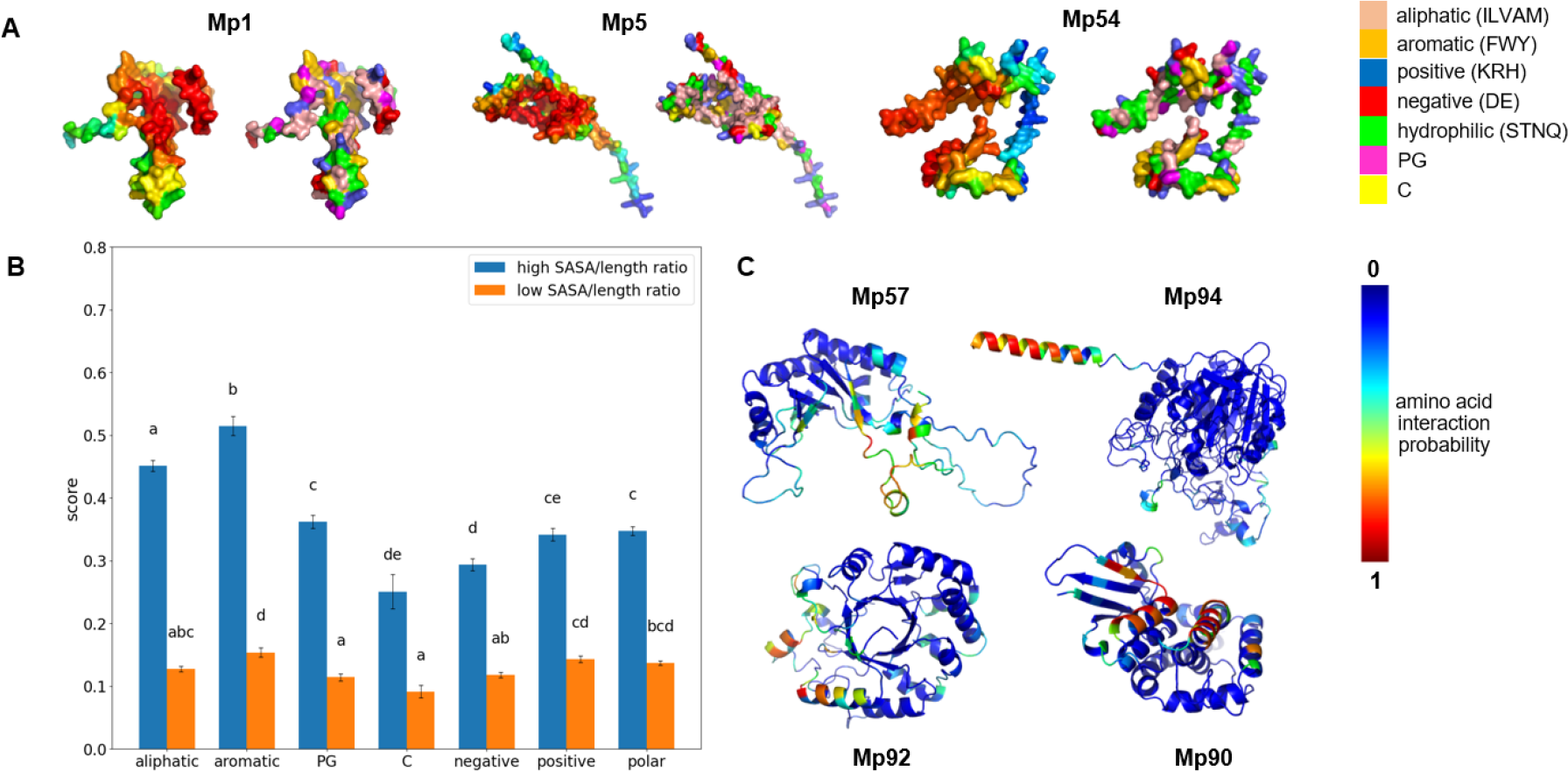
Interactive properties of *M. persicae* salivary effector candidate proteins predicted using ScanNet. **A**, examples of proteins in the disordered category, shown in surface representation with coloring according to ScanNet predicted amino acid interaction probability (left) or Zappo amino acid side chain chemical property color scheme (right) (see color bars). **B**, average ScanNet predicted amino acid interaction probability for different amino acid types. Each value is the mean *±* SE of >80 amino acid residues; means that do not share a letter are significantly different (p < 0.05, one-way ANOVA with Tukey post test). Data are presented separately for the 36 effector candidates with lowest (orange) or highest (blue) solvent accessible surface area (SASA)/length ratio. **C**, examples of proteins in the ordered category, shown in cartoon representation with coloring according to ScanNet predicted amino acid interaction probability (see color bar).

Topsy-Turvy 3D (TT3D) is a recently developed deep learning model demonstrating state-of-the-art performance in predicting protein-protein interaction networks at large scale with good generalizability across species (Sledzieski *et al*., 2023; Singh *et al*., 2022). We used TT3D to predict interaction probability between *M. persicae* effector candidates and proteins in the *Arabidopsis thaliana* proteome. GO term enrichment was analysed for plant proteins predicted as likely to interact with an effector candidate. In this set of plant proteins, some types of GO terms were strongly enriched, including terms associated with response to toxins, endomembrane system, cytoskeleton regulation and redox (Supplementary Fig. 3). This suggests that *M. persicae* salivary effectors might manipulate transport processes that are fundamental to the functioning of the host plant cell, and reactive oxygen species (ROS).

## Discussion

The increasing availability of data about proteins in herbivorous insect pest saliva provides an exciting opportunity to perform large-scale computational analyses to gain overall insights into protein structure, function and evolution. In this study, we applied state-of-the-art computational biology methods to predict the properties of a salivary effector set of a phloem-feeding herbivore and major agricultural pest, the green peach aphid Myzus persicae. We decided to base our analyses on protein structure information, taking advantage of the recent revolution that has occurred in protein structure prediction, and the clear benefits of including structure information in function prediction and detection of evolutionary relationships (Barrio-Hernandez *et al*., 2023; Derbyshire & Raffaele, 2023; Gligorijević *et al*., 2021; Jumper *et al*., 2021; Ruidong *et al*., 2022; Seong & Krasileva, 2023). We predicted structures for 71 effector candidate proteins, and structure served as input information for prediction of function and interaction properties.

### Properties of the *M. persicae* salivary effector repertoire predicted

Confidence in the accuracy of predicted structures arose from the very close agreement between two structure prediction algorithms, AlphaFold and OmegaFold. Both algorithms are proven to be highly accurate despite fundamentally different methodology. For almost all proteins of interest, there was no major disagreement between the predictions. For relatively disordered proteins, the algorithms generally predicted different conformations with a similar proportion of random coil and secondary structure, and proteins with well-defined tertiary structures were readily superimposable with very low RMSD. Importantly, in subsequent analyses using structures or sequences to predict functional and interactive properties, we included proteins predicted as predominantly disordered this subset includes proteins with known effector activity, such as Mp1, MpC002, Mp42, Mp55, Mp57, Mp58 and Mp64 (Atamian *et al*., 2013; Bos *et al*., 2010; Elzinga *et al*., 2014; Liu *et al*., 2022; Mugford *et al*., 2016; Pitino *et al*., 2011; Rodriguez *et al*., 2017; Thorpe *et al*., 2018). We observed clusters of related proteins among the effector candidates, and some relationships were detected based on either sequence or structure similarity, but not both. If sequence similarity only is detected, the proteins may be disordered with little defined structure and share a common ancestor; if structure similarity only is detected, it is possible that convergent evolution has occurred due to favourability of a certain structure for effector function, or the proteins share common ancestry and the sequences have diverged while structure is conserved (Seong & Krasileva, 2023).

Function prediction and comparison to a dataset of randomly chosen *M. persicae* proteins resulted in discovery of protein activities enriched among salivary effectors: protein interaction, carbohydrate degradation, signalling and small molecule interaction. Protein interaction is indeed an expected activity of a plant pest effector, and is essential for targeting of host environment processes as well as for activation of effector-triggered-immunity upon recognition by Nucleotide-binding Leucine-rich Repeat proteins (NLRs) (Ngou *et al*., 2022). For example, *M. persicae* effectors Mp1 and Mp64 were found to interact with host proteins in *A. thaliana* and *N. benthamiana*, resulting in alteration of host processes and enhanced aphid fecundity by largely unknown mechanisms (Liu *et al*., 2022; Rodriguez *et al*., 2017). Predicted protein interaction activity is enriched among the relatively disordered effector candidates only, which emphasizes the importance of studying these proteins. The structure and other properties of these disordered effector candidates may be poorly predicted due to lack of existing knowledge, and yet this set of proteins is likely to harbour the most interesting effector activities that manipulate the host plant environment. Among the predicted enzymatic functions within our effector candidate set, carbohydrate degradation is common and may be involved in digestion/feeding, degradation of the plant cell wall, or perhaps suppression of host defences through binding carbohydrates with elicitor activity (e.g. chitin). Other enzymatic activities were also detected for individual effector candidates, including several redox activities, which may process ROS in order to suppress the plant immune response (Goggin & Fischer, 2021). Signalling activity may be powerful in interfering with the host immune response, while lipid or small molecule binding activity could be involved in interaction with the plant cell membrane, signalling or enzyme catalysis. Attempt to predict plant host target proteins using TT3D suggested activities enriched among targets include intracellular transport, especially membrane trafficking, and regulation of ROS. It is well established that herbivorous insect trigger ROS production and some of their effectors are known to suppress this (reviewed by Wang *et al*., 2023). Moreover, secretory membrane trafficking plays a key role in plant-pathogen interactions (reviewed by Yun *et al*., 2023), and aphid effector Mp1 is known to target Vacuolar Protein Sorting Associated Protein52 (VPS52), a protein involved in membrane trafficking between late endosomes and late Golgi (Rodriguez *et al*., 2017).

### Limitations of this study

The most fundamental limitation of this study resides in the underlying -omics data: some effector candidates may be contaminants in proteomics data, effector identification pipelines may miss detection of effectors for example due to the use of artificial diet collections, and amino acid sequences may be incorrect due to the challenges associated with gene model prediction. Another important limitation is the lack of knowledge of insect salivary effector proteins in general. These proteins are understudied, probably fast-evolving and there is a lack of representation of similar proteins in databases. This leads to unpredictability of the proteins, as predictions mainly rely on artificial intelligence trained on current knowledge and available models. It is possible that some of the disordered structure observed in predictions is incorrect, arising from lack of representation of similar proteins in the PDB. Moreover, while disordered structure might point to important roles in virulence and plant immunity, it will be less useful in evolutionary analyses. Although it has been shown that structure prediction enables more sensitive detection of distant evolutionary relationships between proteins, this is presumably only the case for proteins featuring regions with defined tertiary structure (Barrio-Hernandez *et al*., 2023; Seong & Krasileva, 2023). For proteins with disorder spanning the majority of the primary sequence, it is likely that sequence would be much more highly conserved than structure, and structure prediction might not lead to more accurate prediction of properties, or perhaps even introduce noise that decreases accuracy.

### Future perspective

Our study highlights that predicting the properties of a phloem-feeding insect salivary effector repertoire based on a subset of proteins presents a significant challenge. As mentioned above, the *M. persicae* effector repertoire set is unlikely complete due to challenges associated with effector identification, and structural prediction did not unveil predicted functions for nearly half of the 71 effector candidates used in this study.

In addition, our work highlights the essential requirement for experimental approaches to study aphid effector structure and function. Critically, the identification of host targets, especially for intrinsically disordered effector candidates, will be necessary to provide the opportunity to explore the structures of effectors and their targets in complex and unveil potential mechanisms of action. The combination of experimental and computational structural biology approaches to characterize effector repertoires, and their targets, in less-studied organisms, promises to be a powerful tool. For example, experimentally determined crystal structures of Fusarium oxysporum *f. sp. lycopersici* Avr1 and Avr3 facilitated computational prediction, based on AlphaFold, of a new structural family of effectors (Yu *et al*., 2023). Our work presented here will help guide future experimental and computational work aimed at understanding the virulence activities of aphid effectors, including those predicted as predominantly disordered. With a lack of genetic resistances in both crop and model plants to most aphid species, preventing effector activity may provide an alternative strategy towards crop protection. The development of such strategies will rely on a detailed understanding of how aphid effectors perturb processes in the host environment.

## Methods

### Selection of effector candidates

*M. persicae* salivary effector candidate protein sequences were curated from previously published experimental studies (Bos *et al*., 2010; Elzinga *et al*., 2014; Harmel *et al*., 2008; Liu *et al*., 2022; Thorpe *et al*., 2016; Vandermoten *et al*., 2014) (Supplementary File 1). Only proteins for which the full-length coding sequence of the gene could be found were included. SignalP 6.0 was used to predict the presence of a signal peptide in effector candidate protein sequences, with options ‘Eukarya’ for organism and ‘Slow’ for model mode (Teufel *et al*., 2022). Candidates were excluded if a signal peptide was not predicted to be present in the sequence, or if sequence length was >700 amino acids.

### Selection of effector candidates

The file containing sequences in FASTA format for all predicted proteins of *M. persicae* clone G006 was downloaded from AphidBase (Index of /download /annotation /OGS3.0/ (genouest.org)). 71 proteins were randomly chosen for structure prediction. The file containing details of all entries in the RCSB Protein Data Bank was downloaded (Berman *et al*., 2000). 71 entries were randomly chosen and the structure files (.pdb) downloaded using custom Python code (supplementary data). Only entries consisting of a single protein chain, in which NMR was not the structure solution method, were used.

### Structure prediction

ColabFold was used to run structure prediction algorithms (Mirdita *et al*., 2022). Each protein sequence, with predicted signal peptide removed, was individually submitted to the AlphaFold2_mmseqs2 notebook with adjustable settings specified by the following dictionary: “model_type”: “AlphaFold2-ptm”, “use_templates”: false, “use_amber”: false, “msa_mode”: “MMseqs2 (UniRef+Environmental)”, “num_ models”: 5, “num_recycles”: 3, “num_ensemble”: 1, “pair_mode”: “unpaired+paired”, “stop_at_score”: 100.0, “stop_at_score_below”: 0, “recompile_padding”: 1.0, “recompile_all_models”: false. The rank number 1 model protein structure was used for subsequent analyses. Each protein sequence was individually submitted to the OmegaFold notebook with adjustable settings specified by the following dictionary: “num_cycle”: 4.

### Structure comparison

To quantify structural similarity between AlphaFold and OmegaFold structure predictions, protein structure files (.pdb) were uploaded to the DALI server via the web browser (Dali server (helsinki.fi)) using the ‘Pairwise’ function (Holm *et al*., 2023). To discover homologous proteins based on structure, structure files (.pdb) were submitted to the Foldseek search server via the API using custom Python code (supplementary data), with default search settings (van Kempen *et al*., 2023). To discover clusters of structural similarity among the effector candidates, all the structure files (.pdb) were compared pairwise exhaustively using the Foldseek easy-search command with the following options: –exhaustive-search, –alignment-type 2, -e inf (van Kempen *et al*., 2023).

### Sequence comparison

To discover homologous proteins based on sequence, protein sequences were compared to the UniRef30 database using the HHBlits search server (HHblits | Bioinformatics Toolkit (mpg.de)) with the following search settings: E-value cutoff for inclusion: 0.1, Number of iterations: 8, Min probability in hitlist (%): 20, Max target hits: 10000 (Gabler *et al*., 2020; Mirdita *et al*., 2017; Remmert *et al*., 2011). To discover clusters of sequence similarity among the effector candidates, the multiple sequence alignments output from ColabFold AlphaFold for each protein were compared pairwise exhaustively using the HH-suite3 hhalign command with default options (Steinegger *et al*., 2019).

### Clustering

To cluster proteins based on pairwise similarity score matrices, the Markov clustering (mcl) algorithm was used (v14.137) with inflation factor of 2 (Van Dongen, 2008). Clusters were found after structure-based alignment using Foldseek and selection for protein pairs with e<0.1, or sequence-based alignment using HHalign and selection for protein pairs with e<0.01.

### Structure data processing

Data such as scores assigned to amino acids by algorithms were processed using custom Python code (supplementary data). Solvent accessible surface area (SASA) was calculated using the PDB.SASA.ShrakeRupley module from the Biopython package (Cock *et al*., 2009).

### Function prediction

Protein structure files (.pdb) were uploaded to the DeepFRI server via the web browser (DeepFRI (flatironinstitute.org)) (Gligorijević *et al*., 2021). The output files were processed using custom Python code (supplementary data). For each dataset, abundance of each Gene Ontology (GO) term was calculated by multiplying the number of occurrences of the GO term by the average confidence score.

### Interaction prediction

Protein structure files (.pdb) were uploaded to the ScanNet server via the web browser (ScanNet Webserver (tau.ac.il)), using the ‘protein-protein’ setting (Tubiana *et al*., 2022). *A. thaliana* proteome sequence and structure data was gathered from The Arabidopsis Information Resource (Araport11 protein sequences based on representative gene models) and the AlphaFold Protein Structure Database (reference proteome UP000006548). The Python D-SCRIPT package was used to generate Foldseek 3di sequences for each unique *A. thaliana* protein with a predicted structure available, and to predict probability of interaction between each *M. persicae* salivary effector candidate protein and each *A. thaliana* protein using the Human Topsy-Turvy 3D model (Sledzieski *et al*., 2023; Singh *et al*., 2022; Sledzieski *et al*., 2021). All *A. thaliana* proteins with a predicted score of ≥ 0.9 for interaction with any effector candidate were used for analysis using the GO term enrichment tool provided by The Arabidopsis Information Resource (TAIR).

### Data visualization

PyMOL (The PyMOL Molecular Graphics System, Version 2.0 Schrödinger, LLC) was used to visualize protein structures, and Jalview was used to colour protein structures according to amino acid side chain chemical property (Waterhouse *et al*., 2009). Other data was visualized using custom Python or R code (supplementary data), or GO terms were visualized using GO-Figure with similarity cutoff (-si) equal to 0.1 (Reijnders & Waterhouse, 2021).

### Statistical analysis

Statistical analysis was performed using custom R code (supplementary data) (R Core Team, 2021).

## Data availability

All the data and code underlying this study have been deposited at Zenodo. Data exploration is available in a Streamlit app twaksman001/Mp-Effector-App (github.com).

## Author-Recommended Internet Resources

We recommend the Streamlit app which is convenient for interactive exploration of the data underlying this study. We also recommend PyMOL (PyMOL | pymol.org) for viewing protein structures, and files for this purpose have been deposited at Zenodo.

## Acknowledgements

We acknowledge the Research/Scientific Computing teams at the University of Dundee for providing computational resources and technical support, use of which has contributed to the results reported within this article. This work was supported by the European Union (ERC Consolidator grant, project number 101000997, APHIDTRAP). We also acknowledge Dr. Jade Bleau for input in the selection of effector candidate proteins.

## Author Contributions

T.W., J.I.B.B. and W.N.H. conceived and designed research; T.W., E.A. and S.R.F. performed computational analyses; T.W., E.A. S.R.F., J.IB.B.. and W.N.H. contributed to data interpretation; T.W. and J.B. wrote the manuscript. All authors commented on the manuscript.

