## Supplementary Figures for "Computational prediction of structure, function and interaction of aphid salivary effector proteins"

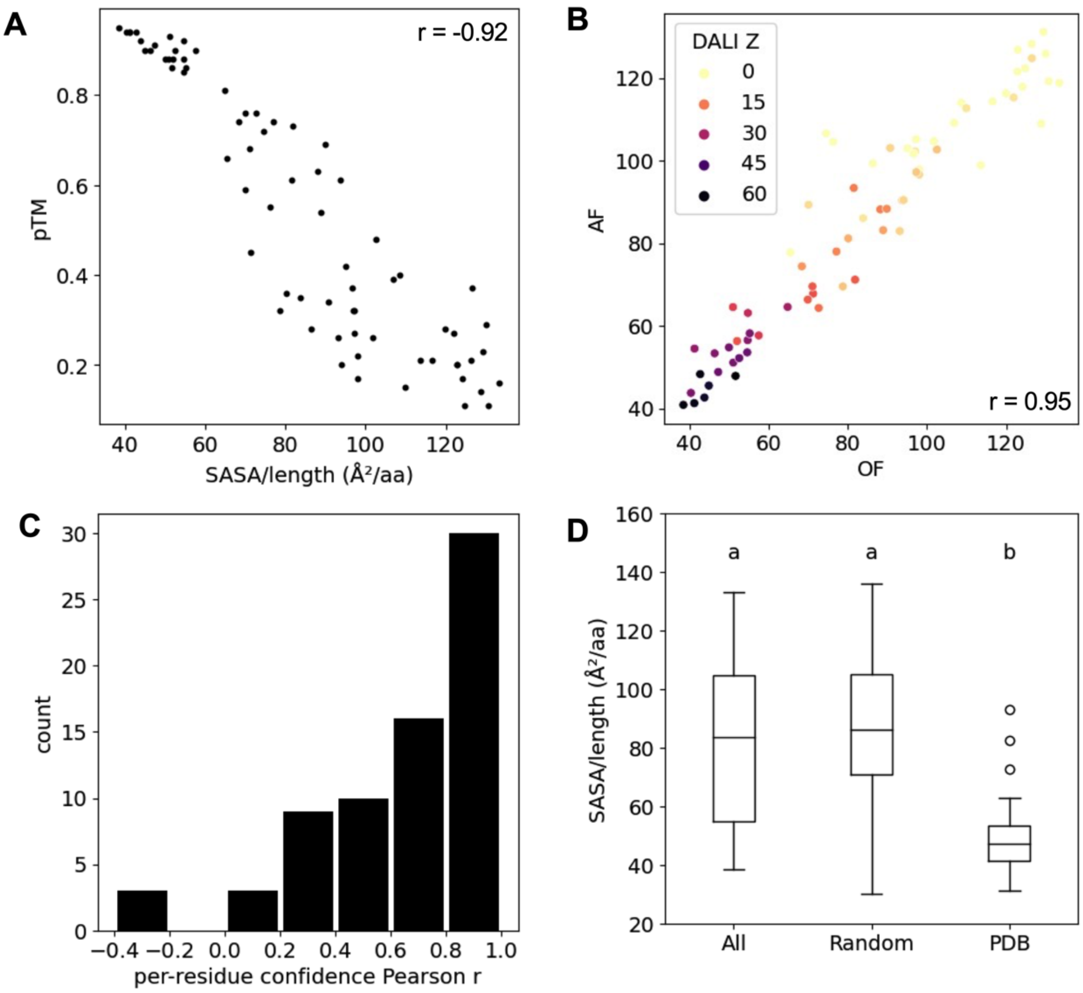


**Supplementary Fig. 1.**
Quantification of *M. persicae* salivary effector candidate protein disorder and similarity of structure prediction between AlphaFold (AF) and OmegaFold (OF). A, predicted template modelling (pTM) score and solvent accessible surface area (SASA)/length ratio of each protein, using the AlphaFold structure prediction. B, SASA/length ratio of AlphaFold and OmegaFold structure predictions for each protein, colored according to DALI Z-score between the pair of structures. C, histogram of Pearson correlation coefficients between AlphaFold and OmegaFold, which were calculated for each protein using the AlphaFold and OmegaFold per-residue confidence score for each amino acid residue in the protein. D, SASA/length ratio of different sets of proteins: all *M. persicae* effector candidates, random *M. persicae* proteins, random proteins from the RCSB Protein Data Bank (PDB). Each value is the mean ± SE of 71 proteins; means that do not share a letter are significantly different ($p < 10^{-14}$, one-way ANOVA with Tukey post test). Pearson correlation coefficient (r) value is indicated if applicable.

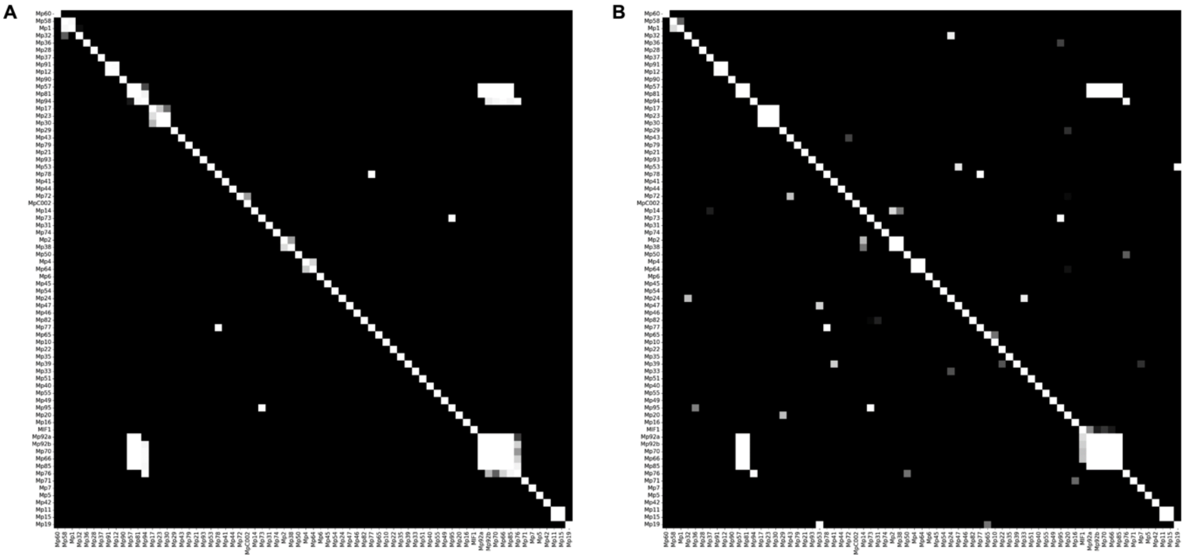

**Supplementary Fig. 2.**
Clusters of *M. persicae* salivary effector candidate proteins based on pairwise alignments. White colour indicates expect value (e)=0. A, Foldseek was used to align all pairs of protein structures, black colour indicates e=0.1. B, HHalign was used to align all pairs of protein sequences, black colour indicates e=0.01.

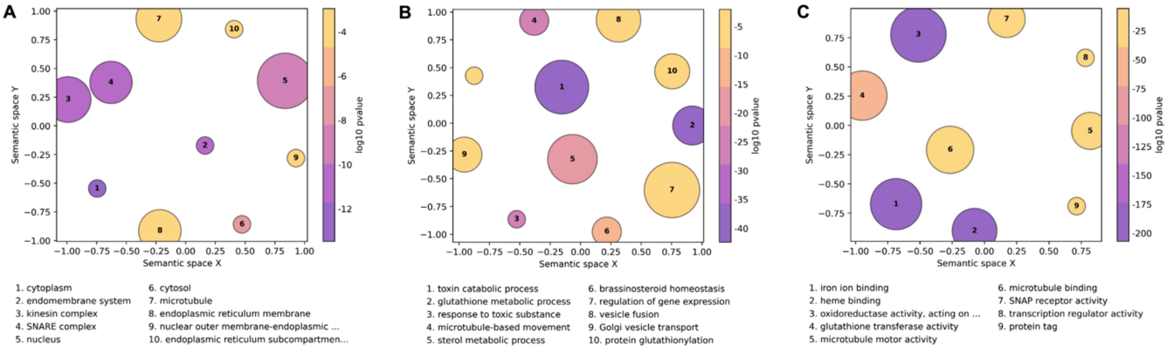


**Supplementary Fig. 3.**
Gene Ontology (GO) terms enriched in *A. thaliana* proteins predicted to interact with *M. persicae* salivary effector candidate proteins. Topsy-Turvy 3D was used to predict probability of interaction between each *M. persicae* effector candidate and each protein in the *A. thaliana* proteome. GO term enrichment of *A. thaliana* proteins with a predicted score of ≥0.9 for interaction with any effector candidate was analysed. Visualization of enriched GO terms was performed using GO-Figure! In the graphs, similar GO terms are close in space, circle size is proportional to number of GO terms in a single group, colour indicates degree of enrichment (p value), and the top 10 enriched GO terms are labelled. A, cellular component GO terms. B, biological process GO terms. C, molecular function GO terms.
